# Lung Epithelial Cells Can Produce Antibodies Participating In Adaptive Humoral Immune Responses

**DOI:** 10.1101/2021.05.13.443498

**Authors:** Erya Gao, Wenwei Shao, Chi Zhang, Meng Yu, Hui Dai, Tianrui Fan, Zhu Zhu, Weiyan Xu, Jing Huang, Youhui Zhang, Zhihai Qin, Xiaoyan Qiu

## Abstract

It is generally believed that the main source of antibodies is B lymphocytes. In this study, our results first revealed that B cell-deficient mice could not produce antibodies specific for TI-Ags; however, mice with B cell deficiency could produce TD-Ag-specific antibodies, although antibody production was delayed compared with that in BALB/c mice after primary TD-Ag challenge. Subsequently, we identified that mouse lung epithelial cells could produce and secrete Ig, including IgM, IgA or IgG, which could display TD-Ag-specific antibody activity. Notably, the production of TD-Ag-specific antibodies by lung epithelial cells was found to be dependent on CD4+ T cells but not CD8+ T cells. Our findings indicate for the first time that B cells are not the only source of TD-Ag-specific antibodies but are essential for rapid TD-Ag-specific antibody production by non-B cells. This discovery may reveal a new mechanism for the production of specific antibodies.

## Main Text

Antibodies, also known as immunoglobulins, have evolved over 200 million years(*1*). They are the most complex and abundant protein family in mammals and are widely distributed in the blood and mucosal tissues(*2*). Although Ig can be classified into five isotypes including IgM, IgG, IgA, IgD and IgE(*3*), each Ig molecule has a unique variable region structure to respond to various antigens, so each individual has a vast Ig library with infinite diversity(*4*). Antibodies are vital components of the immune system for the recognition and response to foreign antigens and self-antigens(*5*). Under physiological conditions, spontaneously produced natural antibodies are widely found in the skin, mucosa and body fluids, such as the serum, to prevent the invasion of pathogenic microorganisms as the first line of immune defense(*2*). Natural antibodies show restricted Ig diversity, low affinity and polyreactivity, and natural antibodies mainly include IgM, IgA and IgG(*6*). In general, natural antibodies are thought to be spontaneously produced by B-1 cells or submucosal plasma cells(*7*).

Under antigen stimulation, the body produces antigen-specific antibodies, which can be divided into two types: thymus-independent antigen (TI-Ag)-specific antibodies and thymus-dependent antigen (TD-Ag)-specific antibodies(*8*). TI-Ags can also be divided into both TI-1-Ags and TI-2-Ags(*9*). TI-1-Ags are microbial ligands of Toll-like receptors (TLRs) that include bacterial lipopolysaccharides (LPS), viral RNAs, and microbial CpG DNA. TI-2-Ags have a highly repetitive structure and can deliver prolonged and persistent signaling to B cells by cross-linking multiple BCRs(*10*). TI-2 antigens include pneumococcal polysaccharides, which can extensively cross-link BCRs on B cells to induce TI-Ag-specific antibody production by B cells in a T cell-independent manner(*11*). TI-Ags can elicit robust antigen-specific IgM and IgG_3_ responses(*12*). Generally, the B cells activated by TI-Ags are in the marginal region of the secondary lymphoid organs and have a limited ability to undergo clonal proliferation but can rapidly differentiate into plasma cells to produce antibodies after antigen challenge(*13*). TI-Ags do not induce a memory immune response(*14*). TD-Ags are antigens with a complex structure and are also important antigens for the immune system. In a TD antigen response, the production of specific antibodies requires the help of activated cognate CD4^+^ T helper cells, which express CD154, the ligand for the costimulatory receptor CD40 expressed on B cells(*15, 16*). TD-Ags can induce rapid B cell proliferation and produce antigen-specific antibody responses(*17*); they can also induce the production of isotype-switched antibodies (IgG, IgA and IgE) and generation of memory B cells(*3*).

To date, B lymphocytes have been considered the only source of Ig, and for a century, studies on the source, structure and function of Ig have been mainly based on the understanding of Ig produced by B cells exerting antibody activity. However, over the past 20 years, growing evidence has demonstrated that Ig, including IgM, IgG and IgA, is widely expressed in non-B lineage cells(*18*), including epithelial cells(*19*), neurons(*19*), germ cells(*20*) and cardiomyocytes(*21*). In particular, malignantly transformed non-B cells can highly express Ig(*22*), which can affect cell or tumor biological activity, such as the promotion of cell proliferation, adhesion, migration and tumor metastasis(*23*). However, to date, few studies have focused on non-B cell-derived Ig, which is involved in humoral immune defense functions. Recent evidence has proven that a variety of epithelial cells can spontaneously produce and secrete Ig, including IgM, IgG, and IgA(*24, 25*), which can bind to a variety of bacterial antigens and bacterial double-stranded DNA to exert natural antibody activity and participate in immune defense in the skin and mucous membranes(*25, 26*). However, whether epithelial cells can produce antigen-specific antibodies has not been discussed.

The lungs are a place for gas exchange. During breathing, lung epithelial cells are continuously exposed to many bacteria, fungi, viruses and other foreign substances. To date, it is believed that lung epithelial cells mainly exert physical and chemical barrier functions. For example, lung epithelial cells form a physical barrier through cilia beating and tight junctions between cells to prevent pathogenic microorganisms from invading. Moreover, lung epithelial cells secrete antibacterial and bactericidal substances, such as antibacterial peptides(*27*), active oxygen molecules and nitric oxide, at the mucosal surface to form a chemical barrier that protects the host from infection by pathogenic microorganisms. However, growing evidence has shown that lung epithelial cells have a powerful antigen recognition system, and all TLRs are expressed in lung epithelial cells(*28, 29*); lung epithelial cells are very sensitive to antigen stimulation and have a strong secretory ability, especially secreting various cytokines, such as TNF, IL-1β, IL-8 and IL-33, and chemokines, such as CXCL9, and recruiting a large number of innate and adaptive immune cells(*30, 31*). In addition, lung epithelial cells also express MHC-II molecules and the costimulatory molecules CD80/CD86, which are necessary for antigen-presenting cells and have antigen-presenting capabilities(*32*), and activate CD4^+^ T cells, especially in the lung epithelium after antigen stimulation(*33*). These cells can transdifferentiate into effective antigen-presenting cells to promote adaptive immune responses. However, to date, in the field of immunology, it is not believed that lung epithelial cells can participate in the immune response by directly secreting antibodies.

This is the first report that B cell-deficient mice can produce antigen-specific antibodies against TD-Ags but not TI-Ags. However, antibody production was delayed in B cell-deficient mice compared with BALB/c mice after primary TD-Ag challenge but not after booster immunization. In addition, we demonstrated that mouse lung epithelial cells could produce and secrete Ig, including IgM, IgA or IgG. More importantly, lung epithelial cell-derived IgG and IgA displayed TD-Ag-specific antibody activity. Moreover, the production of TD-Ag-specific antibodies by lung epithelial cells was found to be dependent on CD4+ T cells. In this study, it was reported for the first time that B cells are not the only source of TD-Ag-specific antibodies but are essential for the rapid production of TD-Ag-specific antibodies in non-B cells. These findings may reveal a novel mechanism for the production of antigen-specific antibodies.

### B cell-deficient mice cannot produce TI-Ag-specific antibodies

To better understand non-B cell responses to TI-Ags, we selected CD19-deficient mice (CD19^−/−^ mice), which lack B-1 cells and have conventional B-2 cells that are poorly responsive to thymus-dependent antigens, as a B cell dysfunction model. Additionally, μMT mice, as a mature B cell-deficient model due to the absence of the transmembrane region of the Igμ chain gene, and severe combined immunodeficiency (SCID) mice, which are a T and B lymphocyte-deficient model, were used; BALB/c mice (wild-type) were used as a positive control (Fig. 1A). We first used 4-hydroxy-3-nitrophenylacetyl (NP)-LPS as a TI-1-Ag to immunize the 4 strains of mice listed above, and the NP-specific antibodies of the IgM and IgG3 isotypes were dynamically detected by ELISA. Markedly, wild-type mice quickly produced NP-specific IgM and IgG3 antibodies beginning on day 3 after challenge with NP-LPS; however, neither NP-specific IgM nor NP-specific IgG3 was found in CD19^−/−^, μMT or SCID mice until 9 days after immunization (Fig. 1B). Our results suggested that non-B cells could not produce TI-1-Ag-specific antibodies, which are B cell-dependent. We also used NP-Ficoll as a TI-2-Ag to challenge these mice and then detected NP-specific antibodies. BALB/c mice produced high titers of IgM and IgG3 antibodies in the serum on the 6th day after immunization with NP-Ficoll. Evidently, no NP-specific antibodies were detected in the serum of μMT mice or SCID mice, which was consistent with the classic immunology theory (Fig. 1C). In contrast to TI-1-Ag stimulation, CD19^−/−^ mice also showed NP-IgM antibody production on the 6th day and NP-IgG3 antibody production on the 9th day after TI-2-Ag stimulation. These results suggested that CD19 deletion does not completely block TI-2-Ag-induced B cell activation.

**Figure 1.**
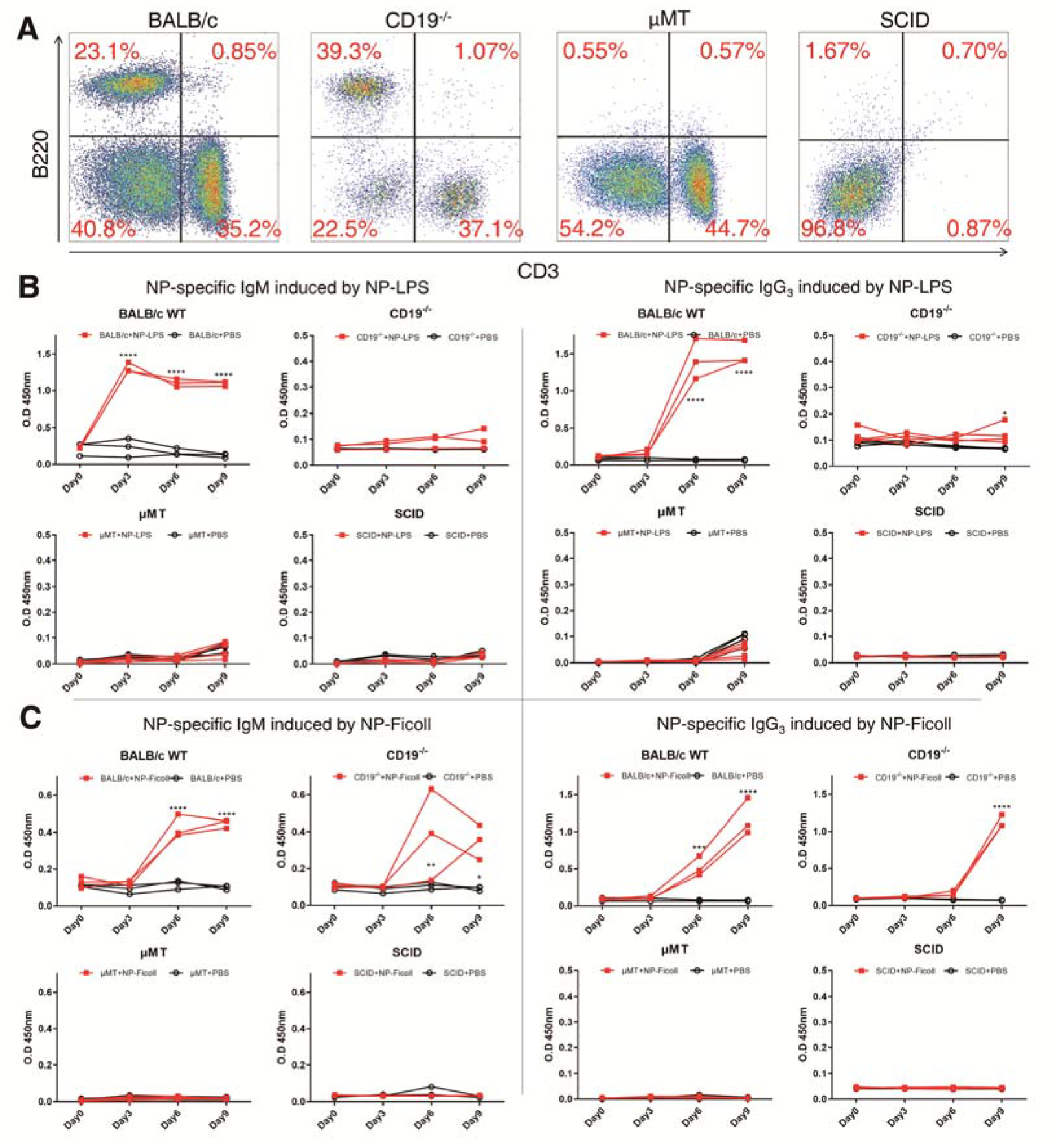
B cell deficient mice could not produce antibodies against TI-Ag. (A) B lymphocytes and T lymphocytes in peripheral blood of BALB/c mice, CD19^−/−^ mice, μMT mice and SCID mice were detected with anti-mouse B220 and anti-mouse CD3 by FACS. (B) The levels of anti-NP-IgM and -IgG_3_ in serum from BALB/c mice, CD19^−/−^ mice, μMT mice and SCID mice after immunized with NP-LPS were detected by ELISA. n=3(C) The levels of anti-NP-IgM and -IgG_3_ in serum from BALB/c mice, CD19^−/−^ mice, μMT mice and SCID mice after immunized with NP-Ficoll were detected by ELISA. n=3 ****P□<□0.0001, ***P□<□0.001, **P□<□0.01, *P□<□0.05, ns P□>□0.05, when compared with the PBS group.

### B cell-deficient mice can produce TD-Ag-specific antibodies, but a lack of B cells or B cell dysfunction results in a significant delay in TD-Ag antibody production after primary TD-Ag challenge but does not affect the rapid production of specific antibodies induced by booster antigen challenge

It has been reported that μMT mice can produce Ig and specific antibodies in response to antigen challenge. To further understand whether non-B cells can produce TD-Ag-specific antibodies and the mechanism of antibody production, in this study, a mixture of NP-KLH (Keyhole Limpet Hemocyanin) and alum adjuvant was injected into the abdominal cavity and foot pad of μMT mice, CD19^−/−^ mice and wild-type mice. Serum was collected from the mice every 7 days after immunization, and NP-specific IgG and total Ig or the number of B cells were detected. As shown in Fig. 2A, immunization with NP-KLH for one week resulted in rapid NP-specific IgG production in wild-type mice but no NP-specific IgG production in either μMT mice or CD19^−/−^ mice.Unexpectedly, from 3 or 4 weeks after immunization, NP-specific IgG was slowly produced at a lower titer in μMT, which is similar to NP-antibody production from 2 or 3 weeks in CD19^−/−^ mice after immunization; in contrast, when NP-KLH was used as a booster immunization 4 weeks after the primary immunization, surprisingly, NP-specific IgG with a similar high titer was quickly detected in wild-type mice and μMT mice. To confirm whether the protein that responded to NP-KLH was NP-specific IgG or another protein, we used an affinity chromatography column by coupling NP-BSA or BSA (as a negative control) with CNBr column material, and the immune serum of μMT mice was incubated with the NP-BSA-affinity chromatography column or BSA-affinity chromatography column. Then, the eluted proteins were analyzed by Western blotting using anti-mouse IgG. The results showed that a 55-kDa IgG heavy chain band was found in the NP-BSA column elution fraction but not in the BSA affinity column elution fraction, indicating that this band was NP-specific IgG (Fig. S1A). In addition, we also determined whether NP-KLH results in an elevated total IgG level or number of B cells in the blood of μMT mice. The results showed that the total IgG level and number of B cells did not change after immunization with NP-KLH (Fig. S1B, C).

**Figure 2.**
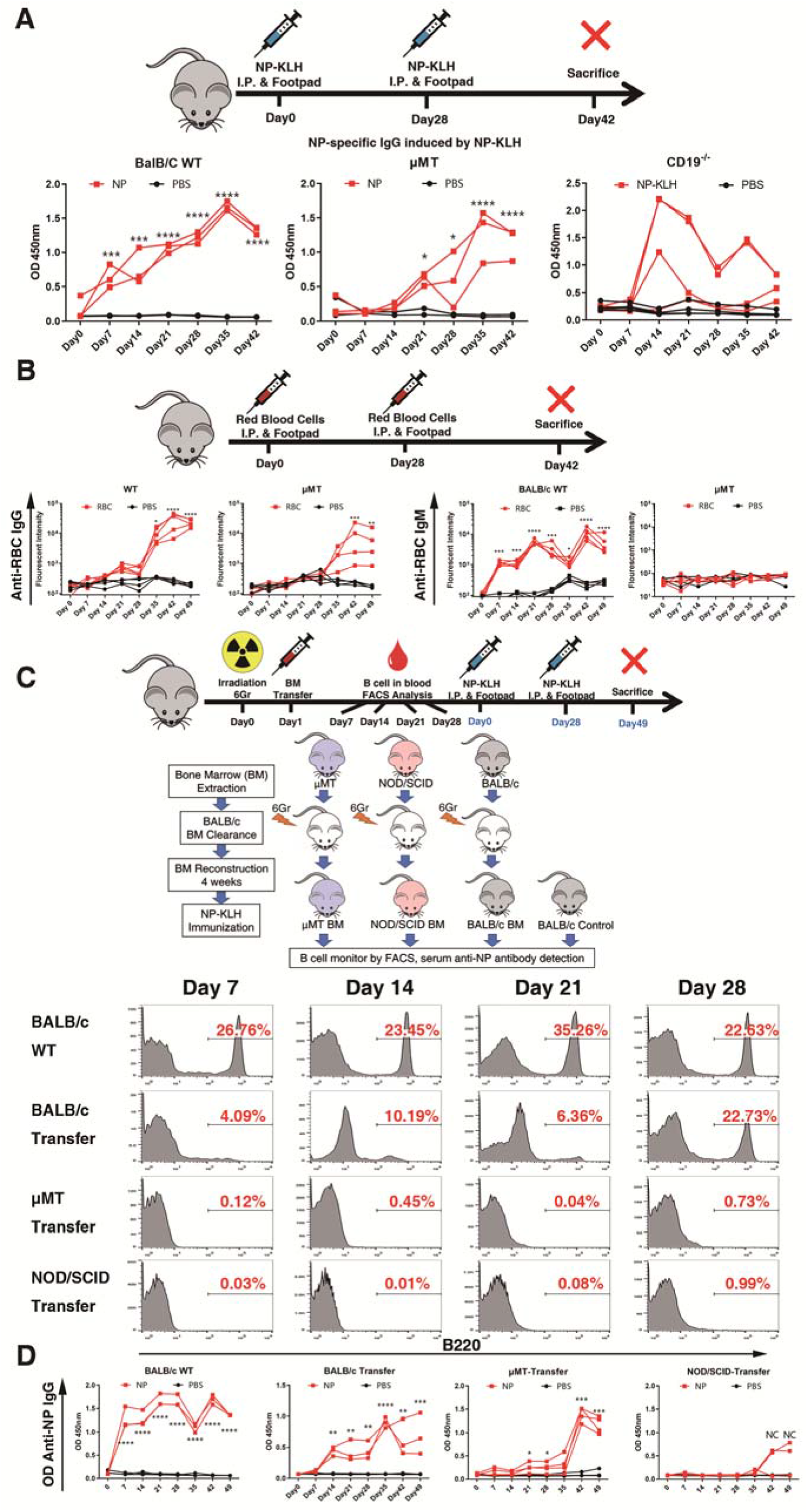
B cell deficient mice could produce antibodies against TD-Ag. (A) We immunized the mice with NP-KLH at day 0, and boosted at day 28. The serum of mice was collected each week until day 42. Then detected the level of anti-NP-IgG in serum from BALB/c mice, μMT mice and CD19^−/−^ mice by ELISA. n=3 (B) The levels of anti-RBC-IgM and -IgG in serum from BALB/c mice and μMT mice after immunized with RBC were detected by ELISA. n=4 (C) The BALB /c mice were first treated with 6 GR radiation, then the bone marrow of μMT and SCID mice were transferred respectively, and the bone marrow transplantation of BALB/c mice were used as positive control. B lymphocytes in peripheral blood of the mice were detected with anti-mouse B220 by FACS. (D) The level of anti-NP-IgG in serum from the mice after immunized with NP-KLH were detected by ELISA. n=4 BALB/c WT mice, the non irradiated BALB/c mice; BALB/c-Transfer, BALB/c mice were adoptive transferred the bone marrow of BALB/c mice; μMT-Transfer, BALB/c mice were adoptive transferred the bone marrow of μMT mice; NOD-SCID-Transfer, BALB/c mice were adoptive transferred the bone marrow of NOD-SCID mice. ****P□<□0.0001, ***P□<□0.001, **P□<□0.01, *P□<□0.05, ns P□>□0.05, when compared with the PBS group.

Next, human RBCs, a low immunogenicity TD-Ag for mice, were also used for primary and booster immunizations in μMT and WT mice. A total of 10^6^ human RBCs were injected into the abdominal cavity, and 4 weeks later, a booster immunization was performed. Specific anti-human RBC antibodies were detected by flow cytometry using human red blood cells as the test antigen. As expected, upon primary immunization with human RBCs, wild-type mice exhibited rapid and high-titer human RBC-specific IgM production, but RBCs slowly induced low-titer human RBC-specific IgG production. However, no human RBC-specific IgM or IgG production was observed in μMT mice within four weeks of the primary immunization. Significantly, after booster immunization, not only wild-type mice but also μMT mice showed rapid and high titers of anti-human RBC-specific IgG. As expected, no RBC-specific IgM was observed in μMT mice (Fig. 2B). The results suggest that the immune memory induced by a humoral immune response to a TD-Ag in μMT mice is B cell-independent. However, under the condition of B cell deficiency, the production of specific antibodies in response to primary immunization was delayed; moreover, the titer was significantly lower in B cell-deficient mice than in wild-type mice (using NP-KLH) or could not be detected (using human red blood cells). These results suggest that B cells play an important role during the first contact with antigens.

To eliminate the difference in genetic background between the above mice, BALB/c mice were first treated with 6-Gy radiation and then transferred with bone marrow from μ MT or SCID mice, and bone marrow transplantation from donor BALB/c mice was used as a positive control. At 4 weeks after transplantation, the bone marrow was basically reconstituted, and the B cells in the peripheral blood of BALB/C recipient mice had recovered to levels similar to those in mice in the untreated group; moreover, no B cells were detected in the peripheral blood of μ MT or SCID bone marrow-transferred mice (Fig. 2C). Then, NP-KLH was used for primary and booster immunizations in the mice. The results showed that a high level of IgG was detected in the nonirradiated BALB/c mice at the first week after immunization, while the level of anti-NP-KLH IgG in the recipient mice adoptively transferred with bone marrow from BALB/c mice gradually increased during the second week after immunization, but the level was significantly lower than that in the untreated mice. The level of anti-NP-KLH IgG in the recipient mice adoptively transferred with μ MT bone marrow increased gradually at 3 weeks after immunization, but it was significantly lower than that in the untreated mice and BALB/c bone marrow-transferred mice. Similar to the previous results, the antibody titer increased sharply after the first week following the booster immunization; however, no specific antibodies were detected in SCID mice (Fig. 2D). These results suggest that TD-Ag specific Ig production in B cell-deficient mice is not caused by the feeding environment or genetic background. Overall, these findings suggest that B cells play an important role in the primary immune response induced by TD-Ag but are not necessary during the response to secondary immunization. In addition, it should be emphasized that under the condition of B cell deficiency, TD antigens can also produce B cell-independent immune memory.

### Ig and TD-Ag-specific antibodies can be expressed and secreted by nonclassical immune cells in the lungs

To search for possible sources of the TD-Ag-specific antibodies in μMT mice immunized with NP-KLH, specific anti-NP IgG and anti-NP IgA were detected in proteins extracted from different tissues by ELISA. The results showed that specific anti-TD-Ag IgG and anti-TD-Ag IgA were widely detected in the liver, kidneys, lungs and small intestine but not in brain tissue (Fig. S2A). When high titers of anti-NP-specific antibodies appeared in the serum, we collected and evaluated mouse lung tissue and the BALF, and NP-specific IgG and IgA in lung tissue and the BALF were detected by ELISA. The results showed that high titers of anti-NP-IgG and anti-NP-IgA were produced in the lung tissue and BALF of μMT mice after immunization (Fig. 3A). Subsequently, the lung tissue was as a model, we explored whether NP-specific antibodies can be express in non B cells under B cell deficiency. The BALF of μMT mice was first collected, and different isotypes of Ig were determined by Western blotting. Our results revealed that the Ig heavy chain, including IgD, IgA, and IgG but not IgM, as well as the light chain, κ and λ, were enriched in the BALF (Fig. 3B). To further identify which local cells in lung tissue can produce anti-TD-Ag antibodies in mice, we analyzed the immune cell subpopulation in the BALF or lung tissue and found a high proportion of alveolar macrophages (AM) in the BALF and high proportions of EpCAM^+^ epithelial cells and lymphocytes in the lung tissue of mice (Fig. S2B).

**Figure 3.**
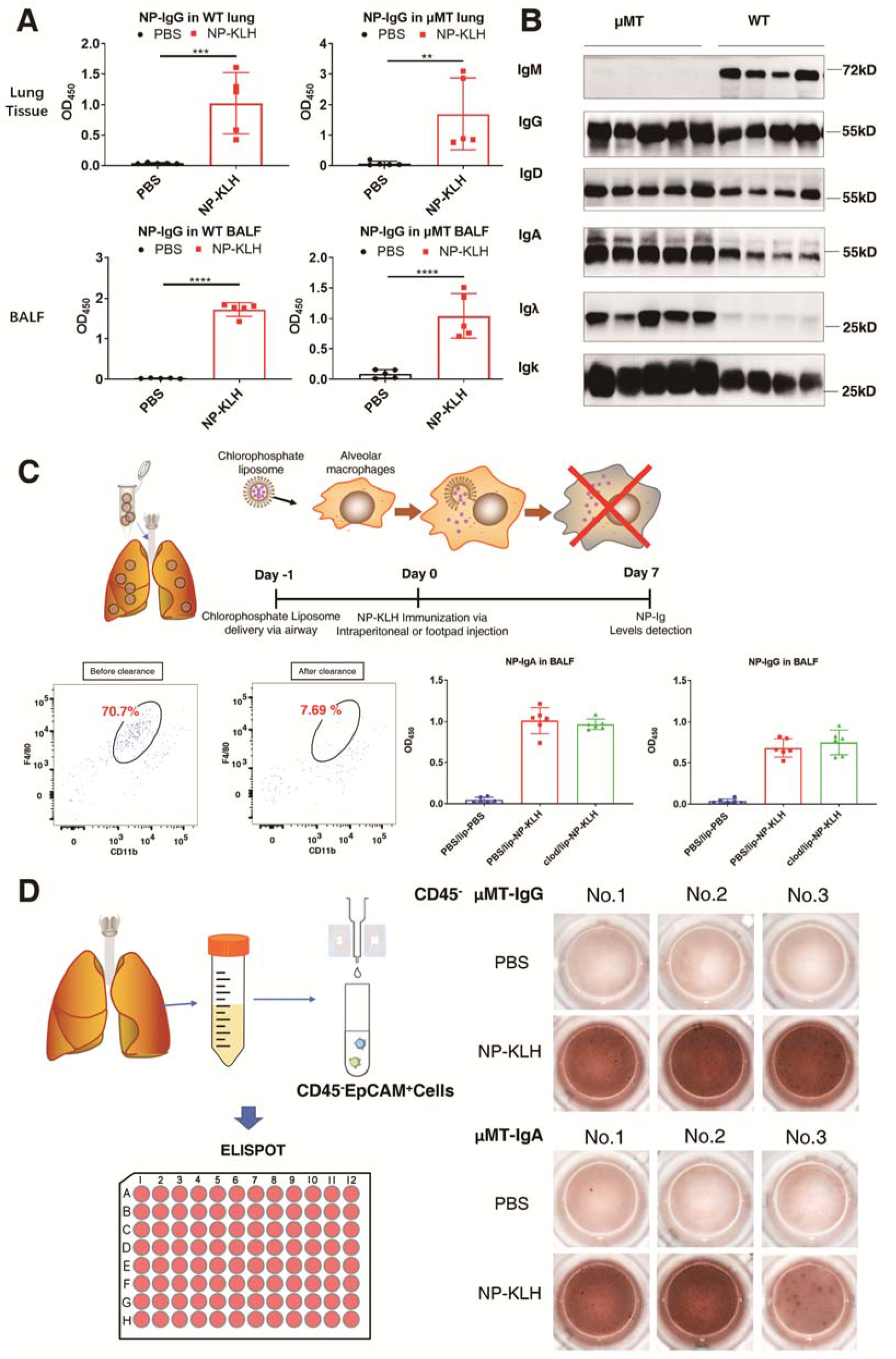
Ig- and TD-Ag-specific antibodies can be expressed and secreted by non classical nonclassical immune cells of in the lungs. (A) The level of anti-NP-IgG in lung tissue and BALF of mice immunized with NP-KLH were detected by ELISA. n=5 (B) The expression of IgG, IgA, IgM, IgD, Igκ and Igλ in BALF of BALB/c mice and μMT mice were detected by western blot. (C) The chlorophosphate liposomes was used to the clearance of alveoli macrophage (AM) of mice. The levels of anti-NP-IgG and anti-NP-IgA in BALF were detected by ELISA. n=6 (D) The secretion levels of anti-NP-IgG and -IgA of lung tissue cells from μMT mice after immunized with NP-KLH were detected by ELISPOT. ****P□<□0.0001, ***P□<□0.001, **P□<□0.01, *P□<□0.05, when compared with the PBS group.

In view of the fact that macrophages have been found to produce Ig(*34*), we next analyzed whether AM can produce anti-TD-Ag antibodies and are involved in the humoral immune response in local lung tissue. To answer this question, chlorophosphate liposomes were used to eliminate macrophages from the mouse lungs by administration via the airways. Clearly, from 24 hours after the administration of chlorophosphate liposomes, the percentage of macrophages in the BALF dropped from 70.7% to 7.69%, which was a significant decrease. Then, the mice were immunized with NP-KLH, and the secretion of specific antibodies into the mouse BALF was detected by ELISA 7 days later. After alveolar macrophage elimination, the production of lung-specific antibodies was not affected (Fig. 3C). This suggests that the local anti-TD-Ag specific antibodies in lung tissue do not depend on AM. To determine whether local nonclassical immune cells can produce anti-TD-Ag specific antibodies, CD45^−^ cells were isolated from the lung tissues of μMT mice immunized with NP-KLH, and the secretion of anti-TD-Ag IgG and anti-TD-Ag IgA was determined by ELISPOT. Clearly, secretion spots indicative of anti-TD-Ag IgG and anti-TD-Ag IgA were identified in the CD45^−^ cells of lung tissues from μ MT mice (Fig. 3D).

### Lung epithelial cells secrete Ig and TD-Ag-specific antibodies

Next, we explored the location of IgA in lung tissue by immunohistochemistry and found that positive IgA staining was mainly observed in bronchial or alveolar epithelial cells of μMT mice and especially after immunization with NP-KLH, the lung epithelial cells was increased in μMT mice (Fig 4A). To further demonstrate that Ig gene rearrangement and transcription occur in lung epithelial cells, we found a mouse lung tissue single-cell sequencing data set (GSE124872) in the GEO database. The data used the Dropseq sequencing method to sequence the whole lung tissue cells of C57BL/6J mice. Since the identification and annotation of lung tissue cell types in these data was previously completed, we directly analyzed the expression of various classes (for heavy chain) or types (for light chain) of Ig in different cells on this basis, and the plasma cell data were used as a positive control. The analysis results showed that Ig was expressed in all types of epithelial cells in the lung tissues and expressed as a cell type-specific marker gene. Type II alveolar epithelial cells (AT2 cells), ciliated cells (ciliated cells) and Clara cells (club cells) mainly expressed Ighm and Igha, goblet cells (goblet cells) mainly expressed Ighm, and plasma cells in lung tissue expressed Ighm, Igha, Ighg2b, and Ighg2c. For the light chain, these epithelial cells only expressed Igk, while plasma cells in lung tissue expressed Igk and Igλ (Fig 4B). These data demonstrated Ig gene rearrangement and transcriptional expression in lung epithelial cells.

**Figure 4.**
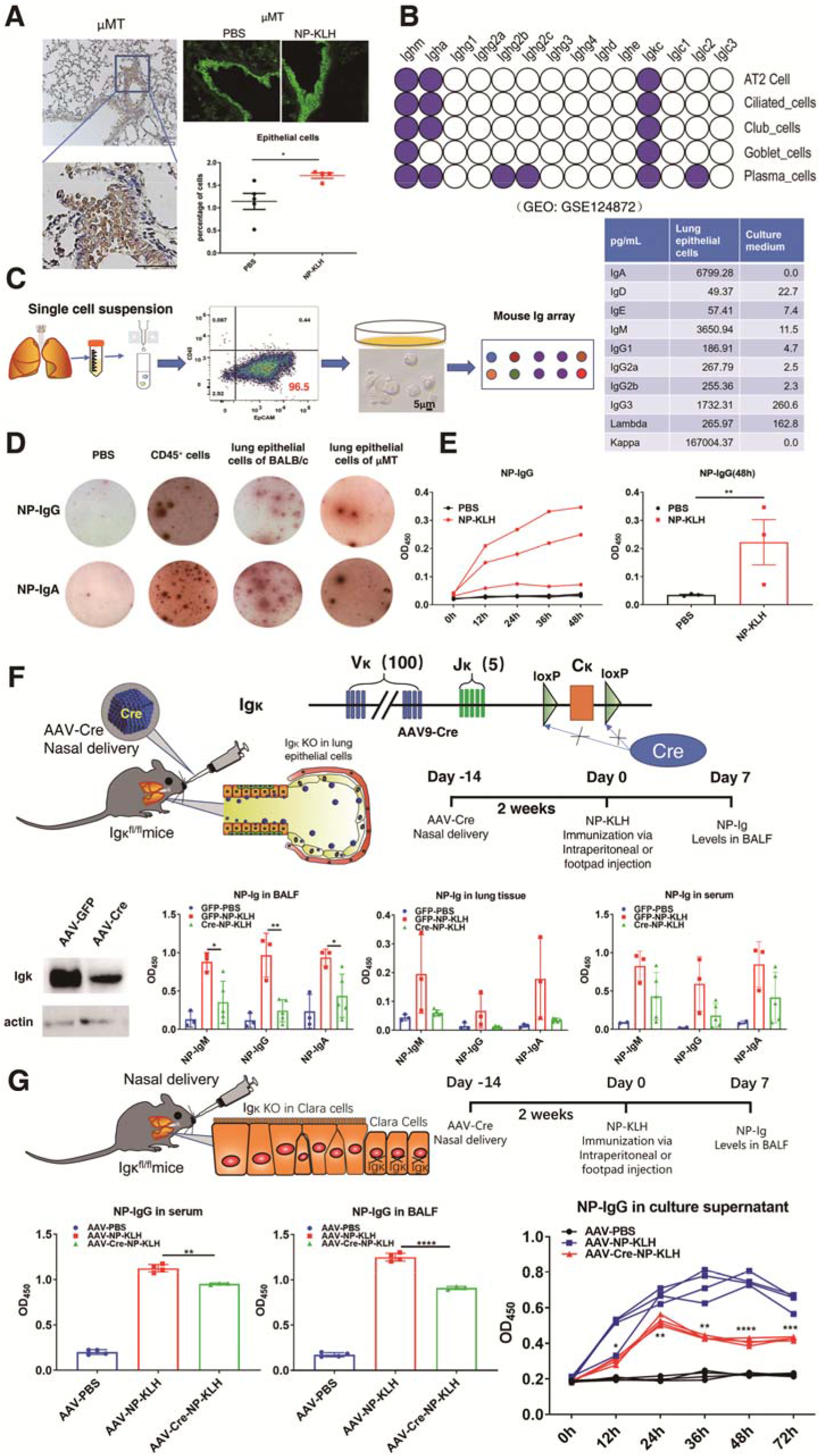
Lung epithelial cells could secrete Ig- and TD-Ag-specific antibodies. (A) The left panel: the local expression of IgA in lung tissue of μ MT mice were detected by immunohistochemical staining. The selected representative image was repeated 3 times independently. Bar=50μm in the figure. The right panel: lung epithelial cells in bronchi or alveoli of μMT mice were detected with anti-mouse CD326 by immunofluorescence (top) and FACS (bottom). (B) The analysis of the various class (for heavy chain) or types (for light chain) of Ig expression in different lung epithelial cells from a set of mouse lung tissue single-cell sequencing data (GSE124872) in the GEO database. (C) The expression of different Igs in the culture supernatant of primary cultured BALB/c mouse lung epithelial cells were detected by the Quantibody^®^ Mouse Ig Isotype Array. (D) The secretion level of anti-NP-IgG and anti-NP-IgA of lung epithelial cells from BALB/c mice and μMT mice after immunized with NP-KLH were detected by ELISPOT. (E) The level of anti-NP-IgG in the culture supernatant of lung epithelial cells (CD45^−^EpCAM^+^) at 0h, 12h, 24, 36h, 48h from μMT mice immunized with NP-KLH were detected by ELISA. n=3, and individual data was represented by each line. (F) We knocked down Igκ in lung epithelial cells by infecting Igc^flox/flox^ mice with AAV-Cre intranasally, then confirmed that AAV-Cre significantly reduced the Igk expression level in lung tissue by Western blot. The levels of anti-NP-IgM, anti-NP-IgA and anti-NP-IgG in BALF, lung tissue and serum of the mice were detected by ELISA.GFP-PBS: n=3, GFP-NP-KLH: n=3, Cre-NP-KLH: n=5 (G) We knocked down Igκ in Clara cells by infecting Igκ^flox/flox^ mice with AAV-CC10-Cre intranasally, the levels of anti-NP-IgG in BALF serum of the mice were detected by ELISA.AAV-PBS: n=4, AAV-NP-KLH: n=4, AAV-Cre-KLH: n=3, and statistical difference between AAV-NP-KLH and AAV-Cre-KLH was analyzed. ****P□<□0.0001, ***P□<□0.001, **P□<□0.01, *P□<□0.05, ns P□>□0.05.

Next, we used magnetic beads to sort CD45^−^EpCAM^+^ lung epithelial cells from both μMT mice and BALB/c mice. Then, the lung epithelial cells of BALB/c mice were cultured, and the different isotypes of Ig in the culture supernatant were detected by ELISA. The results revealed that the epithelial cells cultured in serum-free medium spontaneously secreted high levels of IgM, IgA and Igκ, as well as a certain amount of IgG3, using a Quantibody^®^ mouse Ig isotype array (Fig. 4C). It is worth noting that IgA is the main isotype of Ig present in mucosal tissues. The current consensus in the field of immunology is that this IgA is mainly produced by plasma cells under the mucosa. To confirm that epithelial cells can also secrete IgA, we used the ELISPOT method to detect the secretion of IgA from sorted mouse lung epithelial cells. Lung epithelial cells (CD45^−^EpCAM^+^) sorted with magnetic beads were transferred into an ELISPOT plate at 5×10^5^ cells per well, and sorted CD45^+^ cells were used as a positive control. The results showed that both BALB/c mice and μMT mice contained lung epithelial cells that could express and secrete IgA (Fig. S3), which suggests that under physiological conditions, IgA in the mucosa is partly produced by lung epithelial cells.

To further explore whether specific antibodies can be secreted by lung epithelial cells, CD45^−^EpCAM^+^ lung epithelial cells from both BALB/c mice and μMT mice immunized with NP-KLH were sorted and cultured, and the same number of CD45^+^ cells was used as a positive control. Significantly, by ELISPOT, secreted anti-NP-IgG and anti-NP-IgA were detected (Fig. 4D). We also evaluated anti-NP IgG in the culture supernatant of CD45^−^pCAM^+^ cells from NP-KLH-immunized μMT mice at 0, 12, 24, 36, and 48 h by ELISA. The results showed that the anti-NP-KLH IgG content increased with increasing culture time (Fig 4E). The results indicate that lung epithelial cells can indeed secrete TD-Ag-specific antibodies.

In view of the high expression of Igκ in lung epithelial cells, we first constructed a mouse model of lung epithelial cells with Igκ knocked down. LoxP was inserted upstream and downstream of the mouse Igκ gene constant region. Then, Igκ^flox/flox^ mice were infected with an adeno-associated virus (AAV) containing a Cre enzyme expression vector via nasal drops to knock down Igκ in lung epithelial cells. We first confirmed that AAV-Cre significantly reduced the Igk expression level in lung tissue, and then NP-KLH was used to immunize Igκ^flox/flox^-AAV-Cre mice in the abdominal cavity and foot pads. One week after immunization, the production of specific antibodies was detected in the serum, lung tissue and BALF of the mice by ELISA. The results showed that the levels of specific anti-NP IgM, NP-IgA and NP-IgG were significantly reduced in the BALF after AAV-Cre infection (Fig 4F).

Although AAVs mainly infect epithelial cells, AAV-Cre may still infect other cells in the lung tissue and cause Igκ to be knocked out. Clara cells (club cells) connect the epithelium of the airways and the alveoli and account for approximately 20% of human lung epithelial cells^18^. Considering the large number and strong secretory ability of Clara cells, we decided to target these cells to specifically knock down Igκ in lung epithelial cells. We used AAV-Cre carrying the Clara cell-specific promoter CC10 (AAV-CC10-Cre) and administered this AAV via nasal drops to infect Igκ^flox/flox^ mice and specifically knock out Igκ in Clara cells. Then, NP-KLH was used to immunize Igκ^flox/flox^-AAV-CC10-Cre mice. Similar to the infection with AAV-Cre described above, AAV-CC10-Cre infection resulted in a significant reduction in anti-NP IgG levels in the serum and BALF compared with control treatment (AAV-CC10) one week after immunization. We first confirmed that AAV-Cre significantly reduced the Igk expression level in CC10 expression cells(Fig. S4). Subsequently, CD45^−^EpCAM^+^ cells were further sorted and cultured, and the level of specific antibodies in the culture supernatant was detected by ELISA. Our results showed that compared with the epithelial cells from mice without Igκ knockout, the lung epithelial cells of Igκ^flox/flox^-AAV-CC10-cre mice exhibited significantly reduced specific antibody secretion (Fig 4G).

### Lung epithelial cells can quickly produce TD-Ag-specific antibodies through local airway immunization compared with abdominal cavity plus foot pad immunization

We investigated whether there are differences in the production time and titer of specific antibodies produced by lung epithelial cells between direct airway immunization and abdominal cavity plus foot pad immunization. In addition, to avoid the effects of individual differences in mice, we established a mouse model of unilateral lung immunization. NP-KLH was injected into the left lung of WT mice or μMT mice through a unilateral lung cannula, and the right lung tissue was used as a nonimmune self-control. Moreover, mice immunized with NP-KLH via the abdominal cavity plus foot pad were used as a control for systemic immunity. After immunization with NP-KLH, the production of specific antibodies in the serum was monitored. We first analyzed specific antibodies in the serum of BALB/c mice and found that specific antibodies were clearly identified in the serum following direct unilateral lung immunization or abdominal cavity plus foot pad immunization five days after immunization. Evidently, a higher titer of specific antibodies was achieved in mice by direct unilateral lung immunization than by abdominal cavity plus foot pad immunization (Fig 5A). Then, we analyzed the specific antibodies in the serum of μMT mice and surprisingly found that unlike the mice immunized with NP-KLH via the abdominal cavity plus foot pad, in which NP-specific antibodies were slowly produced for 4 weeks after immunization, the mice immunized via the unilateral lung route quickly produced specific antibodies beginning 12 days after immunization (Fig 5B). The findings indicate that when the lung tissue is directly exposed to NP-KLH, B cells are not required for the response to the initial immunization.

**Figure 5.**
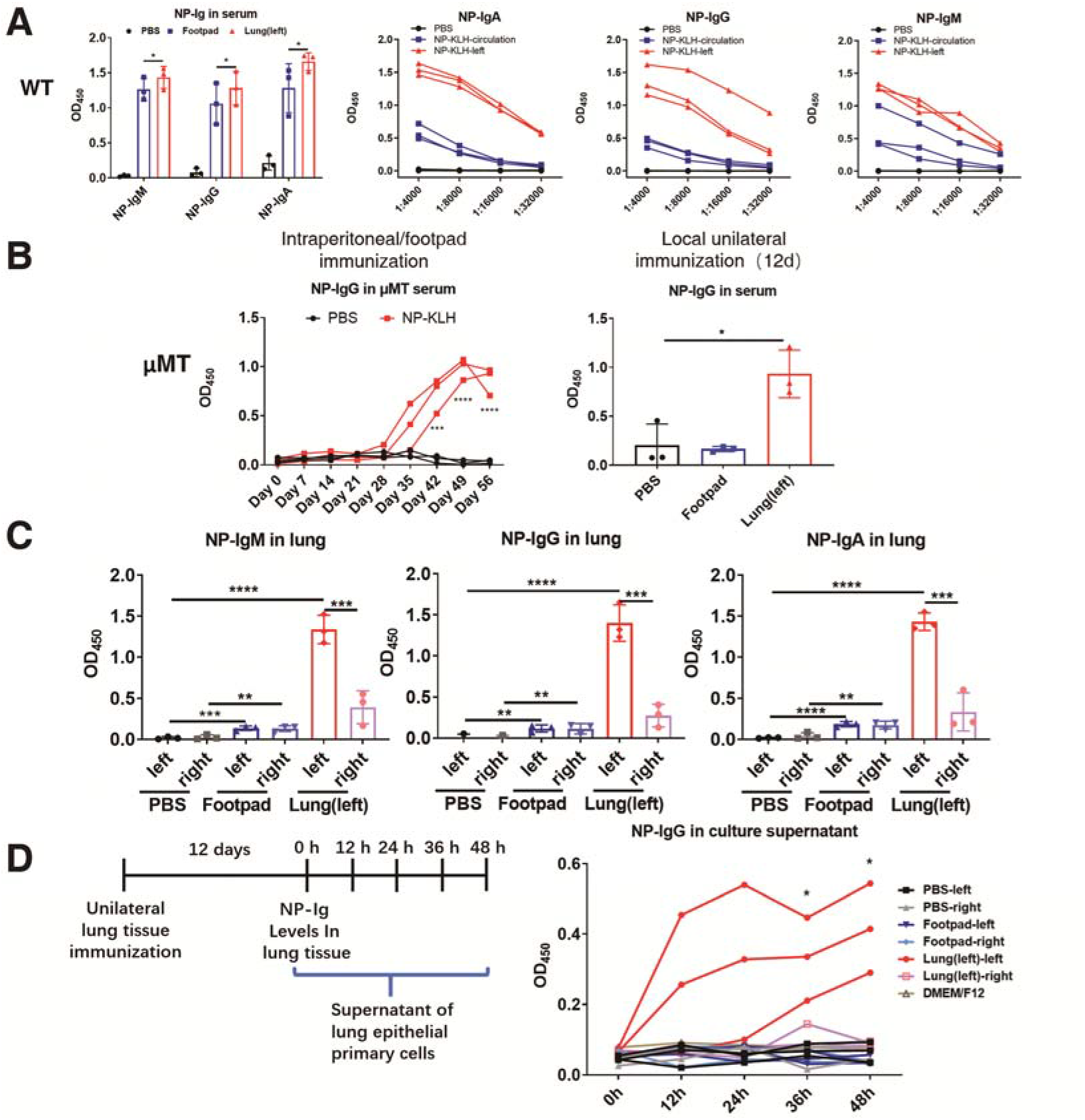
Lung epithelial cells can quickly produce TD-Ag-specific antibodies through the local airway immunization compared with the abdominal cavity plus foot pad immunization. The NP-KLH was injected into the left lung of WT mice or μMT mice through a unilateral lung cannula, the right lung tissue as a non-immune self-control. Moreover, the mice immunized with NP-KLH via abdominal cavity plus foot pad as a control group. (A)The levels of anti-NP-IgM, anti-NP-IgG and anti-NP-IgA in the serum of the BALB/c mice were detected by ELISA. n=3. Statistical difference between NP-KLH-circulation and NP-KLH-left was analyzed. (B) The left panel: the levels of anti-NP-IgG in the serum of the μMT mice immunized with NP-KLH via abdominal cavity plus foot pad were detected by ELISA. n=3. The right panel: The levels of anti-NP-IgG in the serum of the μMT mice were detected by ELISA. n=3. (C) The levels of anti-NP-IgM, anti-NP-IgG and anti-NP-IgA in lung tissue of the BALB/c mice were detected by ELISA. n=3. (D) The levels of anti-NP-IgG in lung epithelial cell culture supernatant of the μMT mice were detected by ELISA. n=3. Statistical difference between Lung (left)-left and Lung (left)-right was analyzed. ****P□<□0.0001, ***P□<□0.001, **P□<□0.01, *P□<□0.05, ns P□>□0.05.

Subsequently, the bilateral lung tissues of unilaterally immunized mice were collected 12 days after immunization with NP-KLH, and the specific antibodies in protein extracted from the bilateral lung tissues of BALB/c mice were detected. Compared to the unimmunized lung tissue, the immunized lung tissue site showed a high titer of NP-specific antibodies, while a lower titer of specific antibodies was found in the right lung tissues of WT mice (Fig 5C). Next, CD45^−^EpCAM^+^ cells from μMT mice treated with left lung immunization or abdominal cavity plus foot pad immunization were sorted 12 days after immunization and cultured, and the culture supernatant was collected at 0, 12, 24, and 48 hours to detect the titer of specific antibodies by ELISA. Similarly, our results showed a high level of specific IgG in the culture supernatant of epithelial cells from the left lung; however, no specific IgG was found in the culture supernatant of epithelial cells from the right lung following left lung immunization, as expected, and no NP-specific antibodies were detected in the culture supernatant of epithelial cells from the bilateral lungs following abdominal cavity plus foot pad immunization (Fig 5D).

To further explore the differences in the expression level and sequence of Ig produced by lung epithelial cells between the immune side and the nonimmune side, we sorted lung epithelial cells (CD45^−^EpCAM^+^) from the tissues of both lungs of mice following unilateral NP-KLH immunization and then performed RNAseq analysis. We first compared the transcriptional levels of different isotypes of Ig in lung epithelial cells from the immune side and the contralateral side of the same mouse. The results showed that the levels of all Ig isotypes, except Igμ, in the lung epithelial cells on the immune side and the contralateral side were significantly increased (Table 1). We also analyzed Ig variable region gene usage and found that several prominent variable region sequences, especially the Igk variable region, were increased on the immune side of the lungs (Table 2).

**Table 1.** Upregulated expression of Ig types induced by NP-KLH unilateral immunization of BALB/c mice (Increase by more than 10 times)

| AccID | Left FPKM | Right FPKM |
| --- | --- | --- |
| Igha | 490.82811 | 172.46123 |
| Ighd | 92.45772 | 30.04517 |
| Ighe | 0.29397 | 0.05649 |
| Ighg1 | 558.79696 | 201.94923 |
| Ighg2b | 119.42246 | 48.05174 |
| Ighg2c | 34.89558 | 9.38191 |
| Ighg3 | 154.27065 | 39.66579 |
| Igk | 0.00908 | 0.00387 |
| Igl | 0.02098 | 0.00806 |

**Table 2.** Usage of NP-KLH inducing the up-regulation and expression of Ig variable region gene V segment in immune side epithelial cells (Increase by more than 3.5 times)

| **AccID** | **Normalized_left** | **Normalized_right** |
| --- | --- | --- |
| **Igkv8-34** | 27.55530005 | 0 |
| **Igkv4-79** | 108.1800669 | 2.939543 |
| **Igkv4-54** | 18.37020004 | 0.979848 |
| **Igkv7-33** | 18.37020004 | 0.979848 |
| **Igkv4-63** | 359.2394674 | 22.5365 |
| **Igkv4-53** | 134.7148003 | 12.73802 |
| **Igkv5-43** | 1492.06847 | 159.7152 |
| **Igkv4-55** | 118.3857336 | 13.71787 |
| **Igkv8-24** | 558.2499677 | 75.44828 |
| **Igkv4-69** | 21.43190004 | 2.939543 |
| **Ighv14-3** | 13320.43616 | 2023.386 |
| **Igkv5-45** | 317.3962339 | 48.99239 |
| **Igkv6-13** | 37.76096674 | 5.879087 |
| **Igkv13-85** | 102.0566669 | 16.65741 |
| **Igkv4-57-1** | 435.7819675 | 77.40798 |
| **Igkv4-70** | 107.1595002 | 20.5768 |
| **Ighv9-1** | 165.3318003 | 44.09315 |
| **Ighv13-1** | 277.5941339 | 74.46843 |
| **Igkv4-61** | 51.02833343 | 13.71787 |
| **Igkv8-21** | 186.7637004 | 52.91178 |

### The production of TD-Ag-specific antibodies depends on CD4^+^ T cells

The characteristic of nude mice is the absence of the thymus, which leads to a lack of T cells. Therefore, we first confirmed the role of T cells in the humoral immune response in nude mice. As expected, a TI-2-Ag could significantly induce TI-2-Ag-specific antibodies; however, when we used a TD-Ag to immunize nude mice, clearly, no NP-specific antibodies were observed in the serum of the nude mice during the whole process of immunization (Fig 6A). The results further indicate that the production of TI-Ag-specific antibodies is dependent on B cells and that the production of TD-Ag-specific antibodies depends on T cells. To further explore whether the production of antibodies by epithelial cells depends on CD4^+^ T cells or CD8^+^ T cells, an anti-mouse CD4 or CD8 monoclonal antibody (mAb) was intraperitoneally injected into mice to eliminate CD4^+^ T cells or CD8^+^ T cells in BALB/c mice, and the same amount of mouse IgG was used in the control group. When no CD4^+^ T cells or CD8^+^ T cells were detected in the lung tissue of mice injected with the anti-mouse CD4 or CD8 mAb, respectively (Fig 6B), the mice were immunized with NP-KLH via abdominal cavity plus foot pad injection. On the 20th day after primary immunization, the production of specific antibodies in the BALF was tested. We found that compared with control treatment, CD4^+^ T cell elimination resulted in no detectable specific antibodies in the BALF, but when CD8^+^ T cells were depleted, antibody production was not affected (Figure 6B), which suggested that the production of TD-Ag-specific antibodies by lung epithelial cells depends on CD4^+^ T cells but not CD8^+^ T cells.

**Figure 6.**
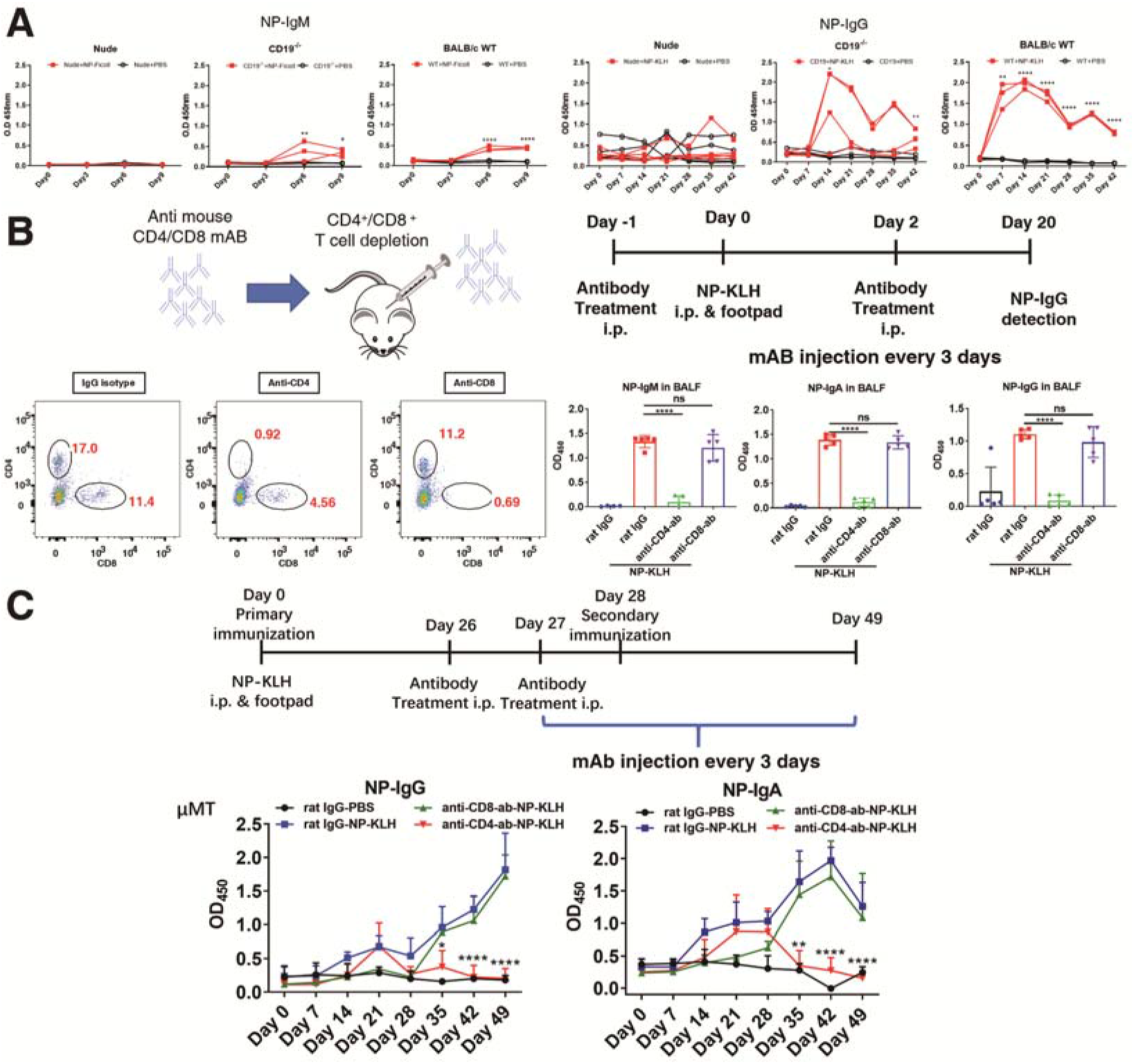
The production of TD-Ag-specific antibodies depends on CD4+ T cells. (A) The left panel: the levels of anti-NP-IgM in the serum of the BALB/c mice, the nude mice and the CD19-/-mice after immunized with NP-ficoll were detected by ELISA. n=3. The right panel: the levels of anti-NP-IgG in the serum of the BALB/c mice, the nude mice and the CD19-/-mice after immunized with NP-KLH were detected by ELISA. n=3. (B) The anti-mouse CD4 or CD8 monoclonal antibody (mAb) was intraperitoneally injected into mice to eliminate CD4+ T cells or CD8+ T cells in BALB/c mice. The left panel: CD4+ T cells and CD8+ T cells in lung tissue of BALB/c mice were detected with anti-mouse CD4 and anti-mouse CD8 by FACS. The right panel: The levels of anti-NP-IgM, anti-NP-IgA and anti-NP-IgG in the BALF of the BALB/c mice after immunized with NP-KLH were detected by ELISA. anti-CD4-Ab, rat anti-mouse CD4 monoclonal antibody; anti-CD8-Ab, rat anti-mouse CD8 monoclonal antibody. n=5 (C) μMT mice were injected with anti-CD4 or anti-CD8 mAb into the abdominal cavity for two consecutive days before the second immunization. The levels of anti-NP-IgG and anti-NP-IgA in the serum were detected by ELISA. anti-CD4-Ab, rat anti-mouse CD4 monoclonal antibody; anti-CD8-Ab, rat anti-mouse CD8 monoclonal antibody. rat IgG-PBS: n=5; rat IgG-NP-KLH: n=5; anti-CD8-Ab-NP-KLH: n=3; anti-CD4-Ab-NP-KLH: n=5. ****P < 0.0001, ***P < 0.001, **P < 0.01, *P□<□0.05, ns P > 0.05.

Next, we explored whether CD4^+^ T cells are important for the maintenance of the immune memory that results in specific antibody production in the lungs of BALB/c mice or μMT mice. First, we immunized BALB/c mice and μMT mice with NP-KLH via the abdominal cavity and foot pads for 4 weeks. Before the second immunization, mice were intraperitoneally injected with an anti-CD4 or anti-CD8 mAb for two consecutive days, and the control group was injected with the same amount of mouse IgG. When no CD4^+^ T cells or CD8^+^ T cells were detected in the blood, the mice were then challenged with NP-KLH, and serum-specific antibodies were monitored. Interestingly, depleting CD4^+^ T cells or CD8^+^ T cells had no significant effect on the production of specific antibodies in the lungs of WT mice (Fig. S5). However, after eliminating CD4^+^ T cells but not CD8^+^ T cells, no specific antibodies were produced in the lungs of μMT mice (Fig 6C). It is suggested that B cells are responsible for the maintenance of humoral immune memory in wild-type mice, but under B cell-deficient conditions, CD4^+^ T cells are responsible for the maintenance of humoral immune memory.

## Discussion

In this study, we demonstrated for the first time that non-B cells could produce specific antibodies and participate in the adaptive immune response. Thus, this finding changed our understanding of the mechanism of antibody production and expanded the classic concept of immunology.

However, some studies have revealed that B cell-deficient mice can produce specific antibodies, such as IgG, IgA and IgE(*35–40*). These findings strongly suggest that specific antibody production is not completely dependent on B cells. Notably, a number of studies have confirmed that non-B cells can widely produce Ig, which has been proven to have natural antibody activity in addition to its important cell biological activities, such as cell proliferation, adhesion, migration and tumor metastasis(*26, 41–44*). For example, IgG and IgA produced by epidermal cells can combine with *Staphylococcus aureus* and *Staphylococcus epidermidis*, respectively, and participate in local innate immunity and epidermal homeostasis(*26*). This evidence further indicates that the previous definition of the origin and function of Ig is limited to the understanding of B-cell-derived Ig, excluding the novel Ig of non-B-cell origin discovered in recent years. However, to date, studies on the function of non-B cell-derived Ig have mainly focused on its cellular biological activity, and its role in the humoral immune response has not yet been explored. Therefore, this study explored the role of non-B cell-derived Ig in the adaptive humoral immune response.

We first used B cell-deficient μMT mice or B cell-dysfunctional mice (CD19^−/−^ mice) and investigated their response to an TI-1-Ag (NP-LPS). The results showed that under conditions of B cell deficiency or dysfunction, mice could not respond to the TI-Ag. These results suggest that the TI-Ag-induced immune response is B cell dependent. Subsequently, μMT mice or CD19^−/−^ mice were used to investigate their responses to TD-Ags (NP-KLH or human red cells), and our observation revealed that B cell deficiency or dysfunction significantly resulted in a delay in specific antibody production following primary antigen stimulation; however, unexpectedly, a rapid production pattern for specific antibodies was found in the context of B cell deficiency or dysfunction following a second antigen stimulation. Obviously, this finding is consistent with previous findings(*38–40*), that is, B cell-deficient mice can produce specific antibodies; however, previous findings did not report that B cells are essential for rapid antibody production after primary stimulation with a TD-Ag antigen but are not necessary for a secondary response.

To further identify the source of specific antibodies in B cell-deficient mice, we extracted protein from the liver, lungs, kidneys, heart, spleen, and brain of mice immunized with NP-KLH and found that in addition to the brain tissue, other tissues showed high levels of anti-NP antibodies, which suggests that most tissue cells may produce specific antibodies. However, we could not exclude the possibility that the high level of anti-NP antibodies in tissues is residual from the circulation; therefore, we focused on observation of the anti-NP antibody level in the BALF. Significantly, a high level of anti-NP antibodies was displayed in the BALF, suggesting that lung epithelial cells or local immune cells may secrete specific antibodies.

The lungs are an organ that communicates with the outside world and are constantly invaded by foreign pathogens; therefore, they have strong immune recognition and host defense mechanisms(*45, 46*); however, whether the lungs can produce specific antibodies is still unclear. Because B cell-deficient mice have been reported to produce TD-Ag-specific antibodies(*36, 38*), in this study, we first identified the source of TD-Ag-specific antibodies under B cell-deficient conditions and found that Ig, specifically IgA, IgG and IgD but not IgM, was abundant in the BALF of μMT mice and that the production of TD-Ag-specific antibodies in the BALF was not dependent on local immune cells, especially alveolar macrophages. Therefore, we used lung epithelial cells as a non-B cell model and systematically explored whether lung epithelial cells can produce TD-Ag-specific antibodies. We first detected whether Ig is present in the BALF of μMT mice and BALB/c mice by Western blotting. Our results clearly showed high levels of IgA, IgG and two light chains of Ig in μMT mice and BALB/c mice; in particular, IgA and Igκ level were significantly elevated in the BALF of μMT mice. Notably, B cell deficiency or IgM heavy chain gene deletion caused a lack of IgM in the BALF of μMT mice, but the mechanism remains to be further explored. Next, to analyze whether the Ig gene was transcribed in lung epithelial cells, we searched a gene expression data set of single lung epithelial cells in the GEO database and found that a variety of single lung epithelial cells showed Ig transcription. Unlike plasma cells that express different classes or isotypes of Ig, lung epithelial cells mainly express Ighm, Igha and Igκ. To identify whether lung epithelial cells can secrete Ig, we sorted and cultured EpCAM^+^ lung epithelial cells from BALB/c mice and found that lung epithelial cells could secrete IgM, IgA, IgG3 and Igκ at high levels. IgA is the main class of Ig in the local mucosa, such as that in the lungs and intestine, which is the first line of defense in mucosal tissue. For a long time, IGA has been considered to be secreted by local plasma cells. The significance of our findings also suggests that not all IgA present in mucous membranes, such as those in lung tissue, is produced by plasma cells; some is secreted by epithelial cells.

Next, we identified whether lung epithelial cell-derived Ig can serve as a TD-Ag-specific antibody. μMT mice or BALB/c mice were immunized with NP-KLH via the abdominal cavity plus foot pads. After booster immunization, we sorted and cultured mouse lung epithelial cells, and the culture supernatant was subjected to ELISPOT and ELISA detection. The results showed that lung epithelial cells could indeed secrete TD-Ag-specific antibodies. To further confirm this finding, we used AAV-Cre, which mainly infects lung epithelial cells, and AAV-CC10-Cre, which targets Clara cells, a population of bronchiolar epithelial cells that accounts for 20% of lung epithelial cells, to knock down Igκ gene expression in Igκ^flox/flox^ mouse lung epithelial cells in vivo and then used NP-KLH to immunize mice via the abdominal cavity plus foot pads. Our results revealed that NP-specific antibody levels in the BALF of Igκ knockdown mice were significantly reduced, especially those of NP-specific antibodies secreted by sorted EpCAM^+^ lung epithelial cells. The results indicate that a long-distance immune pathway can induce the secretion of TD-Ag-specific antibodies in lung epithelial cells.

This led to the question of whether TD-Ag-specific antibodies can be produced in lung epithelial cells by direct TD-Ag challenge in lung tissue. Does direct challenge of lung tissue with a TD-Ag induce quicker production of TD-Ag-specific antibodies? In this study, we developed a unilateral lung immunization technique, with the lung on the unimmunized side acting as a self-control. NP-KLH was used to challenge the left lung of both BALB/c mice and μMT mice via a cannula, and then the level of anti-NP antibodies in the serum was dynamically monitored. After direct lung immunization, specific antibodies appeared in the serum of μMT mice on the twelfth day after the TD-Ag immunization, while at this time point, no specific antibodies were found in the mice immunized via the abdominal cavity and foot pads (generally after 3 weeks of primary immunization); moreover, the antibody titer was the same as that in wild-type mice. It is obvious that local antigen stimulation of the lungs can induce antibody production more quickly under the condition of B-cell deficiency, and the response is independent of B cells when contacting antigens for the first time, which is not consistent with the observations for antibody production in lung epithelial cells induced by remote antigen immunization. We further sorted epithelial cells from the left and right lungs of μMT mice immunized with NP-KLH. As expected, a high level of anti-NP antibodies was observed in the sorted epithelial cells from the left lung but not those from the right lung of μMT mice. In addition, the gene expression profiles of sorted epithelial cells from the left and right lungs of BALB/c mice immunized with NP-KLH were analyzed by next-generation sequencing, and we found that multiple Ig variable region transcripts in epithelial cells from the immune lung were greatly upregulated. Obviously, the specific antibodies in the serum and BALF of mice were derived from non-B cells in the left lung, especially lung epithelial cells.

In view of the facts that induction of TD-Ag-specific antibodies requires CD4^+^ T cell help and that T cell-deficient nude mice cannot produce specific antibodies against a TD antigen, we explored whether the induction of TD-Ag-specific antibodies in epithelial cells also requires CD4^+^ T cell help. We explored the roles of CD4^+^ T cells and CD8^+^ T cells in the production of local lung antibodies by administering anti mouse -CD4 or anti-CD8 mAbs, respectively, and found that depleting CD4^+^ T cells but not CD8^+^ T cells before the first immunization significantly reduced the ability of wild-type mice to produce antibodies but that depleting CD4^+^ T cells just before the second immunization had no effect on the production of antibodies induced by the second immunization in wild-type mice. However, eliminating CD4^+^ T cells but not CD8^+^ T cells significantly abolished the production of specific antibodies in μMT mice. However, the elimination of CD8^+^ T cells had no effect on the production of mouse-specific antibodies. This suggests that the lung epithelial cells of B cell-deficient mice produce specific antibodies that are dependent on CD4^+^ T cells.

CD4^+^ T cells cannot directly recognize natural antigens; they can only recognize antigen peptides bound to MHC II on antigen-presenting cells, such as B cells, DCs, and macrophages. To explore antigen presentation by DCs or macrophages during the immune process induced by NK-KLH challenge, we administered chlorophosphate liposomes to eliminate alveolar macrophages. It was found that macrophage depletion had no effect on the production of specific antibodies in mouse lung tissue. This shows that alveolar macrophages are neither the main antigen-presenting cells for CD4^+^ T cells nor specific antibody-producing cells. Similarly, we used CD11c-DTR mice as a model to discuss whether DCs are involved in antigen presentation by CD4+ T cells; however, when diphtheria toxin was used to eliminate DCs, it decreased the levels of several immune subsets (except for DCs, which were completely depleted), such as T cells and B cells, so the antigen-presenting role of DCs is still uncertain, although depletion did not affect antibody production after treatment with diphtheria toxin for five days (data not shown). In addition, we also paid attention to whether lung epithelial cells may be antigen-presenting cells for CD4^+^ T cells. In 1999, a study found that nasal epithelial cells and airway epithelial cells can take up antigens, further confirming that the airway epithelium has an antigen-presenting function(*47*). Subsequently, scientists discovered that small intestinal epithelial cells, esophageal epithelial cells, corneal epithelial cells and airway epithelial cells all express MHCII molecules and have the ability to present antigens(*32, 48–51*). Among these cells, airway epithelial cells highly express MHCII molecules, and primary cultured lung epithelial cells can present lipid antigens through CD1d molecules(*48*), which plays an important role in the local antigen presentation process. In addition, studies have shown that epithelial cells can also express the costimulatory molecules CD80/CD86 to activate T cells(*52*). Our results confirm that like B cells, lung epithelial cells can produce antibodies. Can lung epithelial cells also function as antigen-presenting cells like B cells? We sorted lung epithelial cells from the immune side and the nonimmune side and then performed RNA-Seq. The results showed that the transcriptional levels of MHCII molecules, such as H2-Ab1 and H2-Eb1, and the costimulatory molecules CD80 and CD86 involved in T cell activation were significantly increased in the lung epithelial cells from the immune side (data not shown). Therefore, our results suggest that lung epithelial cells not only can produce antibodies but also may function as antigen-presenting cells. The detailed mechanism needs to be further explored.

In view of the fact that this study reveals for the first time that lung epithelial cells can produce specific antibodies against a TD-Ag, there are still many scientific issues to be resolved. First, it was established in this study that lung epithelial cells can produce specific antibodies under induction with the TD-Ag NP-KLH and that this process can be completed in B-cell deficient mice, which means that antibody production by lung epithelial cells can be independent of B cells. What is the mechanism by which lung epithelial cells recognize antigens? Second, CD4^+^ T cells are necessary for TD-Ag-specific antibody production by lung epithelial cells. Which subset of CD4^+^ T cells is involved in this event, and which APC is responsible for antigen presentation?

## Methods

### Mice

BALB/c mice, C57BL/6 mice and nude mice were purchased from Charles River Laboratories (Beijing, China). μMT /BALB/c mice were kindly provided by Prof. Qin Zhihai (Institute of Biophysics, Chinese Academy of Sciences). CD19^−/−^ mice were purchased from Model Animal Research Center of Nanjing University (Nanjing, China). Ig _κ_^Flox/Flox^ mice were purchased from Shanghai Model Organisms Center Inc (Shanghai, China). All the animals were bred and maintained in a specific pathogen-free (SPF) conditions in a facility of Department of Immunology, Peking University Health Science Center. All animal studies were reviewed and approved by the Peking University Health Science Center Institutional Animal Care and Use Committee.

### Flow cytometry

To detect B cells, T cells or Macrophage, blood cells were harvest Before staining, cells were blocked with 5% FBS in PBS for 30 min at 4 °C. And then cells were stained with fluorescein conjugated antibodies for 30 min at 4 °C. The isotype control was performed, in which cells were stained with corresponding fluorescein conjugated isotype. Antibodies: anti-mouse IgG-PE, B220-PE CD19-PE-cyA CD3-FITC CD4-PerCP CD8-APC CD45-FITC and CD45-APC were purchased from eBioscience (USA), and CD11b-PerCP and CD138-PE were purchased from Biolegend (USA).

### ELISA

Total IgG, IgM and IgA detection: Mouse IgG, IgA or IgM ELISA Ready-SET-Go kit were purchased from eBioscience (USA). The ELISA plate was coated with capture antibody overnight at 4 °C. After wash with PBST, plate was blocked for 1 hour at RT. Standard sample or 10μg BALF per well were incubated for 1 hour at RT. After wash, detection antibodies were added and incubate at RT for 1 hour. OD450nm was measured with Multiskan FC (Thermo Frisher).

NP specific IgG, IgM and IgA detection: ELISA plate was coated with 4μg/ml NP-BSA (Biosearch) overnight at 4 °C. After wash with PBST, plate was blocked for 1 hour at RT. 100μg lung tissue lysate or 10μg BALF per well was incubated for 1 hour at RT. After wash, detection antibodies were added and incubate at RT for 1 hour. OD450nm was measured with Multiskan FC (Thermo Fisher).

### Mice immunization

50μl, NP-KLH (1μg/μL, Biosearch) was fully mixed with 50μl, alum adjuvant (Thermo Fisher). 8-10 weeks’ mice were immunized by both 80μL of the mixture via intraperitoned injection and 20 μL of the mixture injected to footpads. For unilateral lung immunization, the tracheal of mice was exposed by a 5mm incision, and then 50 μL NP-KLH and QuickAntibody-Mouse2W adjuvant (Biodragon, China) mixture was injected into left lung tissue of mice slowly through a 24G vascular cathete. In the unilateral lung immunization experiment, the amount and type of antigen and adjuvant of the mice immunized via the abdominal cavity and foot pads was the same as the direct lung immunized mice. The mice were then sutured with surgical sutures.

### RBC specific IgG and IgM detection

50μl, Human red blood cells (RBC) separated from human blood samples by density centrifugation were fully mixed with 50μl, PBS. mice aged 8-10 weeks were immunized by intraperitoneal injection of 80μl, of the mixture, with 20μL injected to footpad. Blood samples were taken weekly via outer canthus, serum was isolated by centrifugation. Serum samples were diluted by 2000x with PBS before adding 1μL RBC and gentle mixing by pipetting. After incubation for 30min at 4°C, sample tubes were centrifuged 5000×g for 5 min at 4°C. After discarding supernatant, anti-mouse IgG-PE and anti-mouse IgM-PE-Cy7 antibodies diluted in 100 μL PBS were added to RBC pellet, and mixed by gentle pipetting. After incubation for 30min at 4°C, sample tubes were centrifuged 5000×g for 5 min at 4°C, then washed in 1ml PBS before centrifugation and subsequent re-suspension in 100μL PBS. The sample was then analyzed by FACS, the concentration of RBC-specific IgG or IgM corresponds to relative fluorescent intensity of PE or PE-Cy7.

### Lung and bronchoalveolar lavage fluid (BALF) harvest

After the anesthetization, mice heart was exposed and perfused with sterile PBS for 10 min until the color of lung turning white. The lung tissue was then removed from the mice. For BALF harvest, the trachea of mice was exposed, and an incision was cut under the cricoid cartilage. 1ml sterile PBS was injected into the lung tissue, and pipet up and down for 3 times, and the BALF can be stored at −20°C.

### Protein extraction and Western Blot

Tissues were harvest after perfusion and cut into small pieces. The tissues were lysed with TSD buffer (1% SDS, 50mM pH7.5 Tris-HCl, 5mM DTT and proteinase inhibitor), followed by ultrasonication. Lysate was centrifuged at 10,000rpm for 15 min at 4°C. The supernatant was collected and used as tissue lysate. After protein concentration detection, the reduced tissue lysate by β-mercaptoethanol was separated by SDS-PAGE. For Western blot, the separate proteins were transferred to nitrocellulose membranes. After blocked with tris-buffered saline containing and 5% defatted milk for 2 h at room temperature, membranes were incubated with anti-mouse IgG-HRP, anti-mouse IgM-HRP and anti-mouse IgA-HRP (ThermoFisher), goat anti-mouse Igκ, goat anti-mouse Igλ (Southern Biotech), rat anti-mouse IgD (eBioscince): overnight at 4 °C. Following the wash with TBS, membranes were incubated with the rabbit anti-goat IgG-HRP and rabbit anti-rat IgG-HRP antibodies (Beijing Zsbio. Co.) in the dark. ImageQuant LAS 500 (GE) was used to detect the signal.

### Lung epithelial cell isolation

Lung epithelial cell isolation was performed using gentleMACS dissociator (Miltenyi Biotec). Put lung tissues in the gentleMACS C Tube containing 5ml HEPES buffer, followed by adding 2 mg/ml collagenase IV and 40 U/ml DNase I. Place the C Tube on the dissociator and runs the procedure “m_lung_01”. Remove the C Tube and digest it for 30 minutes at 37°C, 180rpm. Then, place the C Tube back on the dissociator and run the program ‘m_lung_02’. The cell suspension was filtered through a 70μm strainer, and then washed with 5mL HEPES Buffer. The cells were centrifuged at 300g for 10 min and then resuspended with 10ml PBS pH 7.2 buffer containing 2mM EDTA and 0.5% FBS. Cells were centrifuged at 300g for 10 min and repeat this process 2 times. Then the cells were counted and ready for next procedure.

Lung epithelial cells separated by magnetic bead: the anti-mouse CD45 magnetic bead (Miltenyi Biotec) was used to enrich the epithelial cells in single-cell suspension, and then the anti-mouse EpCAM magnetic bead (Miltenyi Biotec) was used to separate the epithelial cells.

For flow cytometry analysis and sorting: stain cells using FITC anti-mouse CD326 (Biolegend), APC anti-mouse CD45 (Biolegend) for 30mins at 4°C. Wash and resuspend cells in 0.5% FBS/PBS, and then flow cytometry sorting can be performed to isolate lung epithelial cells.

For isolate lung epithelial cells, RNA-seq and analysis were performed by NovelBio Bio-Pharm Technology Co., Ltd.

### Lung epithelial cell culture

Primary lung epithelial cells were cultured in DMEM/F12 medium with 0.005 mg/ml insulin, 0.01 mg/ml transferin, 30 nM sodium selenite, 10 nM hydrocortisone, 20ng/ml EGF, and 2%FBS. 2×10^5^ cells per well were seeded in 48-well plate. Before the Ig detection, change the medium into serum-free medium and culture 24h.

### ELISPOT

To detect the secretion of NP specific IgG or IgA in lung epithelial cells, ELISPOT was performed. MultiScreen filter plates were coated with 10μg/ml NP-BSA for overnight at 4 °C. 1× 10^5^ sorted lung epithelial cells were used per well. After the blocking with 10% FBS, cells were added and cultured for 24 h at 37 °C. After washing with PBS, incubate with HRP conjugated detection antibody at room temperature for 1 h. Then following AEC coloration, results were analyzed by ELISPOT reader

### Immunohistochemistry

Lung tissues were fixed in 10% formalin for 72 hours, then embedded in paraffin. The paraffin blocks were sectioned and mounted on slide. After deparaffinization, the paraffin slides were rehydrated using decreasing concentrates of ethanol. Antigen unmasking was performed in pH 9.0 tris-EDTA buffer at a sub-boiling temperature for 5 min. To quench the endogenous peroxidase activity, the slides were treated with 0.3% hydrogen peroxide for 5 min. Then sections were blocked with 10% goat serum for 20 min. Sections were incubated with anti-mouse IgG or IgA antibodies (Bethyl Laboratories, USA) at 37 °C for 1 hour in a humidified chamber. After PBS washing, sections were incubated with HRP-conjugated rabbit anti-goat IgG antibody (Beijing Zsbio. Co.) at 37 °C for 40 min. After washing with PBS, the signal was detected using DAB (Dako, CA, USA). Sections stained without primary antibody were used as negative controls.

### Immunofluorescence on Cryosection

After perfusion, the lung tissue of mice was embedded in OCT, which was to be completely filled into the lung tissue and then the tissue can be stored in liquid nitrogen.

Cut the tissue into slides with a thickness of 5μm. Then the slides were fixed with acetone at room temperature for 5 min, and stored at −30°C. For immunofluorescence, 5% FBS was used to block slides at room temperature for 20 min. Then FITC anti-mouse CD326 (Biolegend) was added and incubated at room temperature for 2 h in the dark. Wash with PBS 3 times, followed by incubating with Hochest33342 at room temperature for 5 minutes. Then mount the slide with 50% glycerin and a cover glass.

### Igκ knocking down in vivo

After anesthesia, Ig κ^Flox/Flox^ mice of 8-10 weeks were treated with 1×10^11^ AAV vectors per mouse by nasal drops. For Igκ knocking down in lung epithelial cells, AAV9-Cre-GFP or AAV9-GFP were used. For Igκ knocking down in lung epithelial cells, AAV9-CC10-Cre or AAV9-CC10 were used. The 20μL droplet were carried out slowly and uniformly to avoid asphyxia.

### Alveolar macrophages depletion in vivo

After anesthesia, the tracheal of mice was exposed by a 5mm incision, and then 50 μL chlorophosphate liposomes (5mg/ml) or PBS was injected into lung of mice slowly and uniformly. 24 hours later, immunization was performed.

### T cell depletion in vivo

In vivo depletion of CD4+ and CD8+ T cell was performed using anti-CD8 (clone 2.43) or anti-CD4 (clone GK1.5) blocking antibodies. Each mouse was injected i.p. with 0.5 mg monoclone antibodies or normal rat IgG (Beijing Zsbio. Co.) in 200 μL PBS two days before immunization, and repeat every three days after that.

### Statistical analysis

All statistical analysis was performed using GraphPad Prism 7.0 software. Data are presented as mean± SEM. For all multiple comparisons, data were compared using 2-way ANOVA. For the data from single comparisons, it was evaluated by the Student’s t test. P value of <0.05 between different groups was considered significant.

## Acknowledgements

We wish to thank Prof. Dalong Ma (Peking University) and Prof. Bo Huang (Chinese Academy of Medical Sciences, in Beijing) for their help and support.

## Funding

This work was supported by grants of XYQ:

National Natural Science Foundation of China (No. 91642206, No. 81971465),

Novel coronavirus research and development plan of Ministry of science and technology (No. 2020YFA0707801),

Non-profit Central Research Institute Fund of Chinese Academy of Medical Sciences (2019PT320006),

Research Institute Fund of NHC Key Laboratory of Medical Immunology, Peking University (BMU2018JDJS010),

Special Projects for Strengthening Basic Research of Peking University (BMU2018JC003).

## Author contributions

X.Q. and E.G. designed the experiment; E.G., C.Z., W.S. and M.Y. contributed to in vitro and in vivo experiments; T.F., J.H. and H.D. contributed to in vitro experiments; Z.Q., and W.X. supplied mice used in this study;; E.G., W.S., C.Z. and Z.Z. carried out the data analysis; X.Q., W.S., C.Z., E.G. and Y.Z. wrote the paper; X.Q. supervised the work as PI. All authors reviewed the manuscript.

