## supplementary material for "Lung Epithelial Cells Can Produce Antibodies Participating In Adaptive Humoral Immune Responses"

**This PDF file includes:**

Figs. S1 to S5

**Fig. S1**

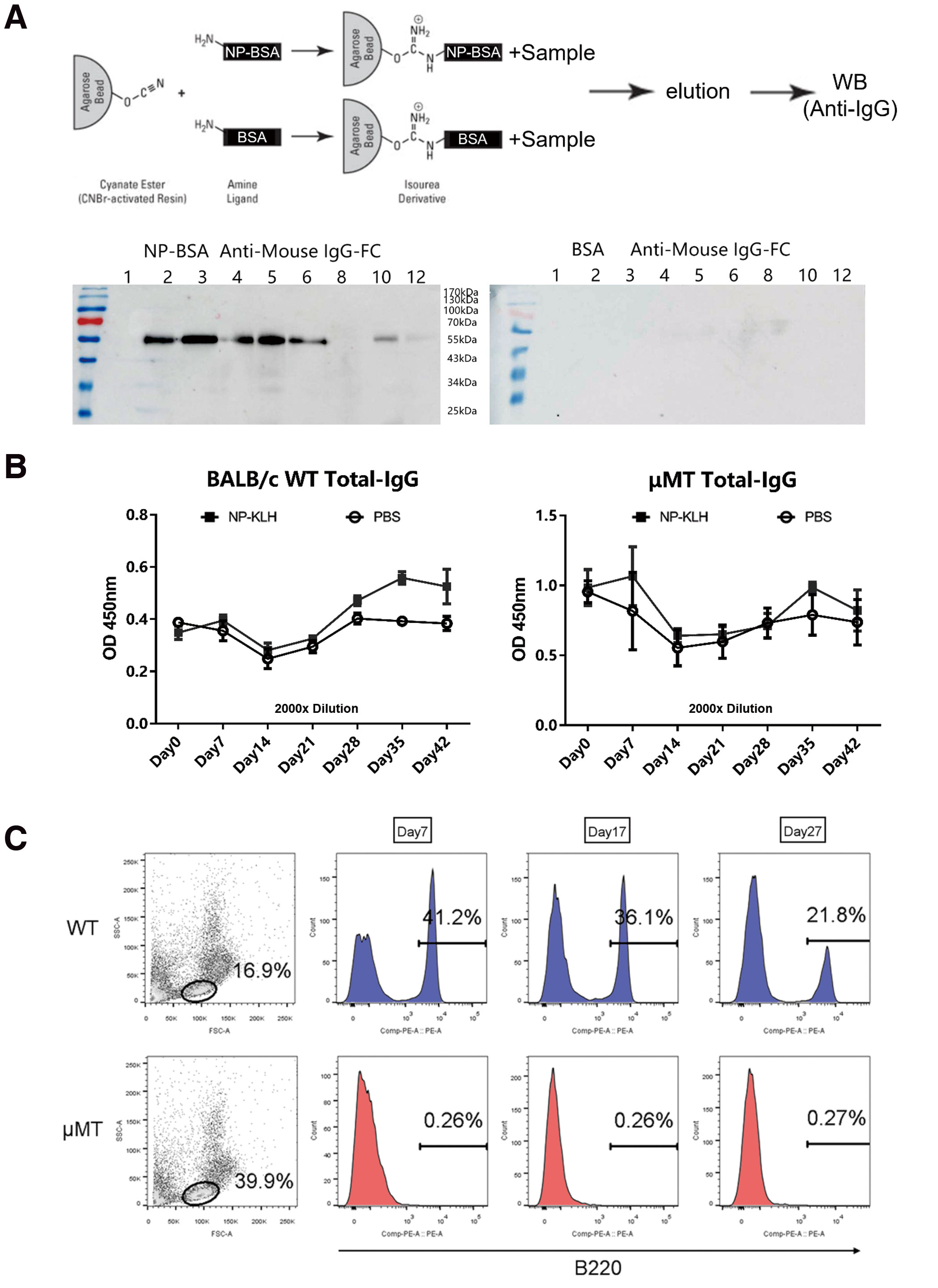

**Figure S1. The positive response detected by ELISA is NP-specific IgG**

（A）We made affinity chromatography column by coupling NP-BSA and BSA with CNBr column material respectively, and the immunized serum of μMT mice was used to incubate and combine with NP-BSA- or BSA-affinity chromatography column, respectively. Then the level of 55 kDa IgG heavy chain band in the eluting components was detected by Western blot. (B) The level of total IgG in serum from the BALB/c mice and μMT mice after immunized with NP-KLH were detected by ELISA. The serum was diluted 2000x n=3 (C) B lymphocytes in peripheral blood of the BALB/c mice and μMT mice were detected with anti-mouse B220 by FACS.

**Fig. S2**

**Figure S2. Lung epithelial cells may express Ig.**

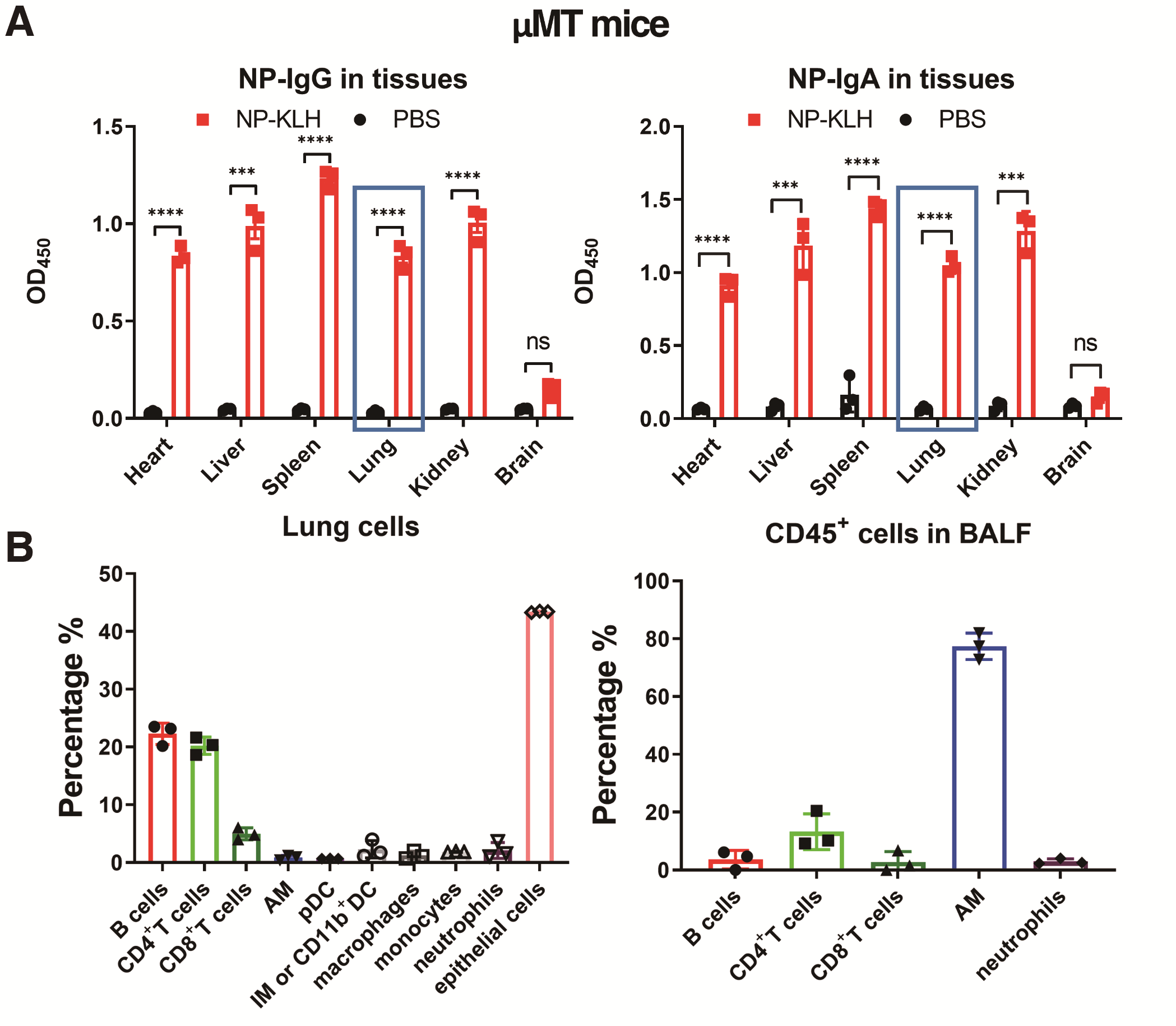

(A) The level of anti-NP-IgG and anti-NP-IgA in various tissues from μMT mice after immunized with NP-KLH were detected by ELISA. n=3 (B) the immune cell subpopulation in the lung tissue and BALF were detected by FACS. n=3. ****P < 0.0001, ***P < 0.001, **P < 0.01, ns P > 0.05.

**Fig. S3**

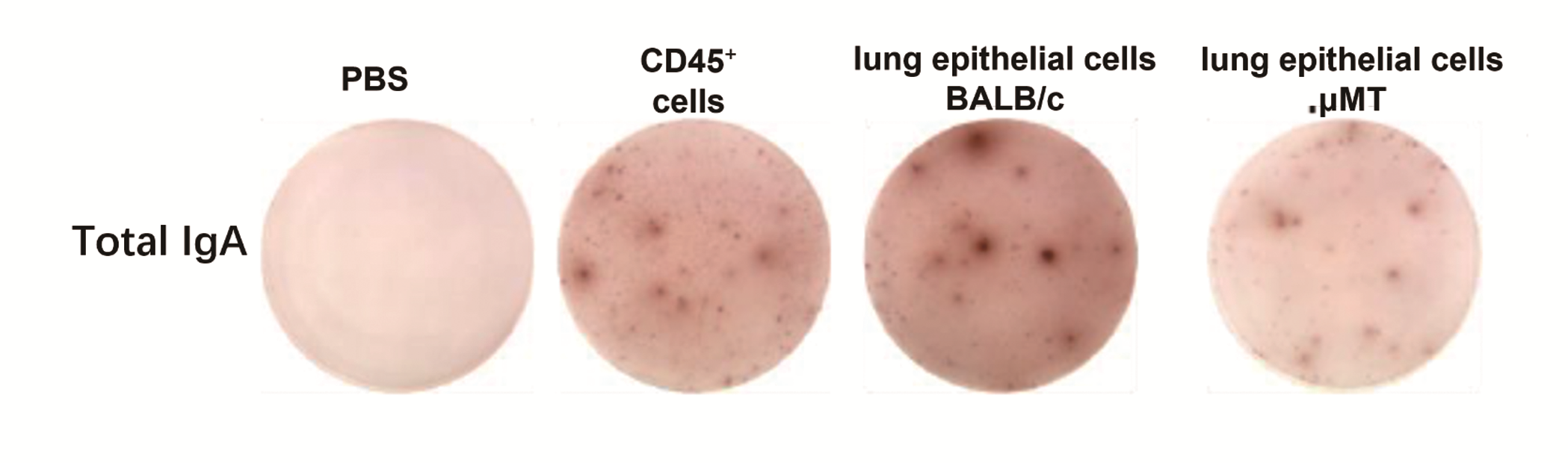

**Figure S3. Lung epithelial cells could secrete Ig- and TD-Ag-specific antibody**

The secretion level of IgA of lung epithelial cells from BALB/c mice and μMT mice after immunized with NP-KLH were detected by ELISPOT.

**Fig. S4**

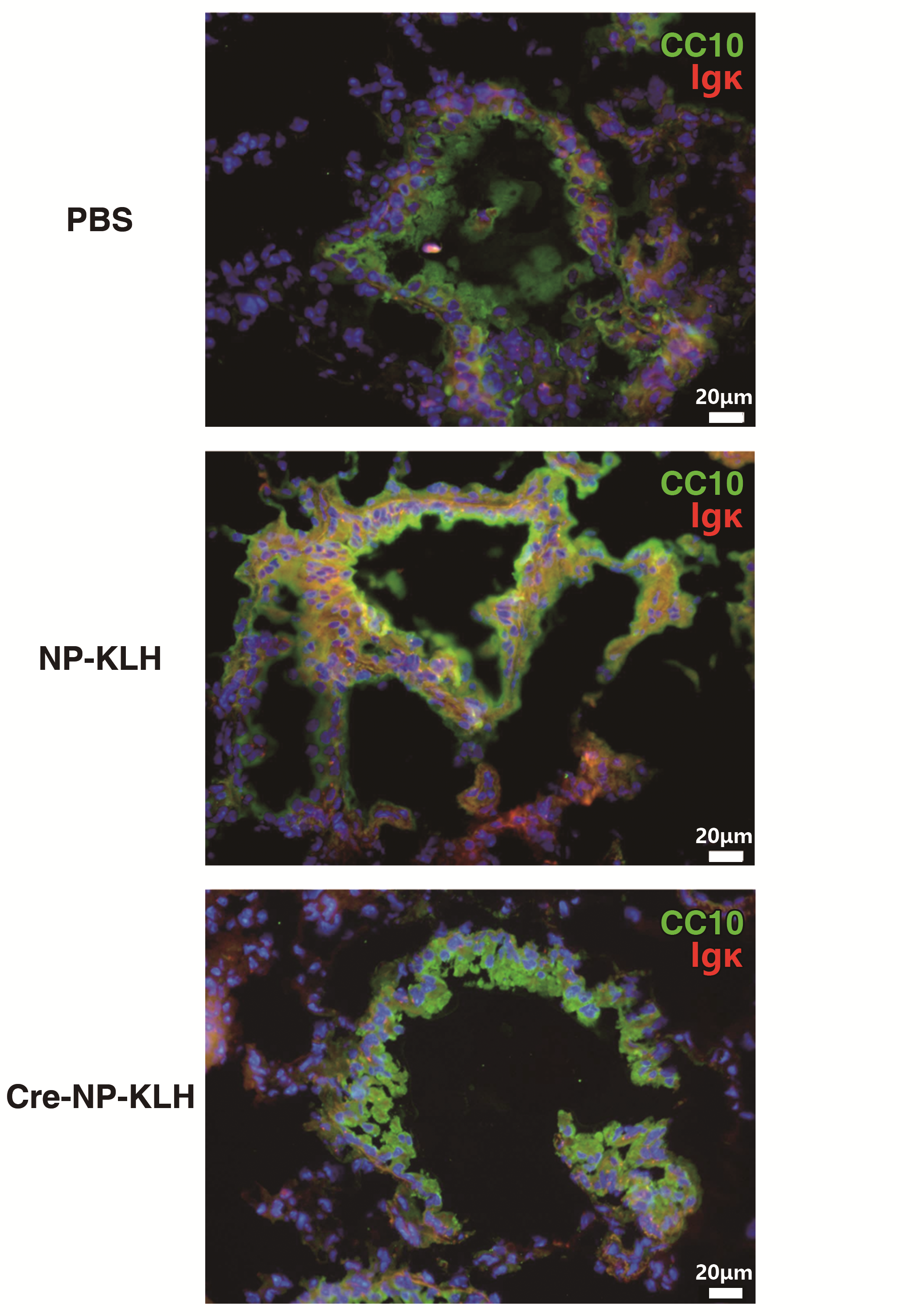

**Figure S4 The Igk expression level in CC10 expression cells of lung**

The protein level of Igκ(red) in CC10 expression cells(green) in lung of Igκ^flox/flox^ mice were detected with anti-mouse uteroglobin and anti-mouse Igκ by immunofluorescence. Bar=20μm.

**Fig. S5**

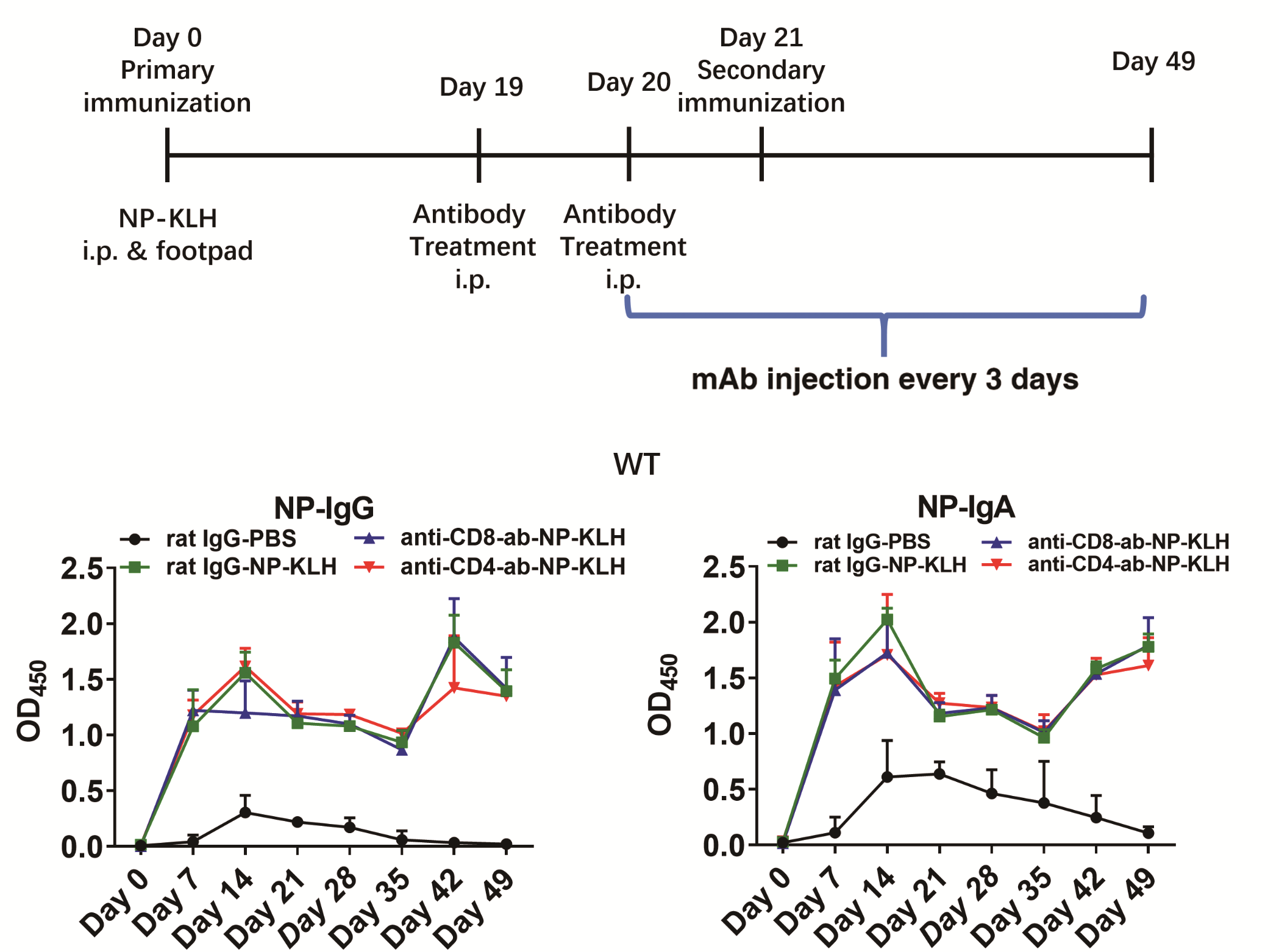

**Figure S5. The maintenance of the immune memory that results in TD-Ag-specific antibodies production did not depend on CD4^+^ or CD8^+^ T cells in BALB/c mice**

BALB/c mice were injected with anti-CD4 or anti-CD8 mAb into the abdominal cavity for two consecutive days before the second immunization. The levels of anti-NP-IgG and anti-NP-IgA in the serum were detected by ELISA. anti-CD4-Ab, rat anti-mouse CD4 monoclonal antibody; anti-CD8-Ab, rat anti-mouse CD8 monoclonal antibody. rat IgG-PBS: n=4; rat IgG-NP-KLH: n=3; anti-CD8-Ab-NP-KLH: n=3; anti-CD4-Ab-NP-KLH: n=4.
